# Short-term stabilities of 21 amino acids in dried blood spots

**DOI:** 10.1101/196295

**Authors:** Jun Han, Rehan Higgins, Mark D. Lim, Karen Lin, Juncong Yang, Christoph H. Borchers

## Abstract

**BACKGROUND:** Dried blood spots (DBSs) have potential use in remote health applications for individual and population diagnosis, and can enable epidemiological surveillance for known and unknown diseases. The preparation and transportation of DBSs from remote settings often exposes these cards to extreme environmental stress that may impact the quality of the diagnostic data. Given these risks, it is essential to investigate the individual stabilities of biomarkers in DBSs. This paper details the stability of routinely-analyzed amino acids (AAs) on DBSs under environmental conditions that simulate a global health workflow.

**METHODS:** The extractions of 21 AAs from three sets of DBSs prepared on cellulose and cotton filter paper were optimized for quantitation by dansylation-UPLC/MRM-MS. The effects of sunlight exposure, temperature, humidity, and storage time were studied.

**RESULTS:** The AAs were stable in DBSs after 4-hour sunlight exposure, and after storage at -20 and 4 °C for 30 days. At 25 and 40 °C, only 7 AAs showed significant concentration decreases over time, while 2 showed concentration increases. The changes were accelerated by high humidity. Histidine was the least stable AA under the conditions tested.

**CONCLUSIONS:** This study provides quantitative data on the short-term stabilities of 21 AAs in DBSs on cellulose and cotton-based filter paper, under environmental conditions that simulate a global-health workflow. These results highlight the importance of assessing the stability of clinically-relevant biomarkers in DBSs. Based on the measured stabilities, we recommend that higher-temperature and high-humidity storage of DBS samples be avoided for AA analysis in remote health applications.

## Introduction

Dried blood spots (DBSs) are often prepared by blotting and drying microliter volumes of capillary blood -- obtained by pricking the heel, finger, or toe --onto filter paper as a simple sampling format. Compared to other sample collection methods such as venipuncture, DBS sampling is less invasive and the samples typically require fewer resources for shipping and storage (1, 2). DBS specimens have been used to screen newborns for inborn errors of metabolism (IEMs) since its first introduction more than five decades ago (3). Because of its simple collection, low sample volume, and ease of transportation, DBS sampling also holds promise for expanding from newborn screening in a hospital setting to enabling the diagnosis of other diseases, particularly from remote settings where blood collection and transportation may be challenging. One example is the use of DBS specimens to enable remote diagnosis of HIV viral load to assess efficacy of treatment (4), an approach recommended by the World Health Organization (WHO) for resource-limited settings. In recent years, there has also been an increased interest in the use of DBSs as a cost-effective replacement for blood plasma or serum in studies involving proteomics (5-7) or metabolomics (8-10), as well as in the community of pharmaceutical research and development for therapeutic drug monitoring, pharmacokinetics (PK), or toxicokinetics (TK) (11).

WHO-recommended and US-Food and Drug Administration (FDA) approved DBS collection cards, considered as medical devices for blood specimen collection, include Whatman 903 Protein Saver and Whatman FTA DMPK-A, B, C, and D cards, all of which utilize cellulose as the absorbing matrix and are distributed by GE Healthcare (Piscataway, NJ, USA), as well as PerkinElmer 226 (previously known as Ahlstrom grade 226) Spot Saver cards, which are manufactured from cotton linter by PerkinElmer (Greenville, SC) (12), etc. Whatman 903 cards are cleared by the FDA for newborn screening for IEMs, and the DMPK-A, B, C, and D cards are often used in PK/TK studies. Of the four types of FTA DMPK cards, DMPK-A and B cards are chemically treated with proprietary reagents that, on contact, induce cell lysis, protein denaturation, enzyme inactivation, and inhibition of bacterial growth. The Whatman 903 and DMPK-C cellulose cards and the PerkinElmer 226 cotton cards are not treated with any chemicals, so there are no introduced chemicals which might interfere with the analysis of low-molecular weight compounds in blood. In an inter-laboratory study, the performance properties of filter paper for whole blood collection between Whatman 903 and PerkinElmer 226 were compared by evaluating the standard substance spiking recoveries of 26 newborn screening analytes, including 5 amino acids (AAs). The results demonstrated the similarities in analyte recovery between the two types of filter paper, indicating that the two types of collection cards provide comparable quantitation accuracies for newborn screening (13).

Most global health programs aim to extend access to health care services for populations residing in settings that are often poorly resourced and distant from a sophisticated diagnostic laboratory. The concentrations and compositions of free AAs in blood have long been implicated as an important indicator of protein adequacy in children malnutrition (14), but nutrition surveys are hard to conduct in global health settings. A recent milestone study revealed that stunted growth in children with malnutrition is associated with low concentrations of all 9 circulating essential AAs (*His, Ile, Leu, Lys, Met, Phe, Thr, Trp, Val*), together with low concentrations of conditionally essential AAs (*Arg, Gly, Glu*) and non-essential AAs (*Asn, Glu, Ser*) in the blood serum (15). DBS sampling offers great potential for remote health applications as a cost-effective sample format for large-scale screening and diagnostic purposes, but the DBS samples often need to be transported from remote sampling locations (e.g., tropical or developing countries) to analytical laboratories where the quantitative measurements of different metabolic or disease biomarkers can be carried out. During sample transportation and storage, DBS-preserved specimens are often exposed to extreme environmental conditions (e.g., sunlight exposure, high temperature, or high humidity) that could significantly affect the chemical stabilities of the different biomarkers, reducing the reliability of any quantitative result. Given these risks, it is therefore imperative to investigate the possible degradation of different biomarkers such as AAs in DBSs under normal and some extreme environmental conditions. In a previous study, the long-term stabilities of 10 AAs, together with a few carnitines, on DBS samples collected from infants and stored at room temperature for a time period of up to 15 years were evaluated using flow injection analysis (FIA)-tandem mass spectrometry (MS/MS),a rapid analytical technique widely used in clinical laboratories for routine screening for IEMs (16). For short-term stabilities of AAs, Golbahar, et al. used a FIA-MS/MS method with neutral loss scanning for AAs and with parent ion scanning for carnitines to check the stabilities of 7 AAs and 10 carnitines in DBS specimens under several testing conditions, including low humidity (<30% relative humidity, RH) and high humidity (>70% RH), and low temperature (up to 4 °C), room temperature (25 °C) and high temperature (37 and 45 °C) for up to 8 days of DBS storage (17). Thus, there have been very few systematic investigations of the short-term stabilities of other AAs in DBSs to date, and little is known about the potential effects of different absorbing matrices -- for example, cellulose versus cotton filter paper-- on the short-term stabilities of different AAs under the various environmental conditions that are typically encountered in a global health workflow.

The purpose of this study was to evaluate the short-term stabilities of 21 routinely-analyzed AAs in DBS specimens prepared on three sets of WHO-recommended and FDA-cleared cellulose or cotton filter paper-based collection cards (i.e., Whatman 903, Whatman FTA DMPK-C, and PerkinElmer 226) under four sets of well-controlled stability-testing conditions. Experiments included (1) one exposure to 4 hours of sunlight irradiation; (2) storage at 20 °C for 2 days, 40 °C for 2 days and then back to -20 °C for 2 more days (three cycles = 144 hours total) under reduced humidity conditions (using silica gel); (3) storage at four temperatures (-20, 4, 25, and 40 °C), with measurements at 2, 6, 15, and 30 days for each temperature; and (4) storage at 75% relative humidity (RH) at 25 or 40 °C, with measurements at 2, 6, 15, and 30 days. To provide precise and accurate quantitation of the AAs in the tested samples, we first improved the sample preparation for AA extraction and combined it with pre-analytical dansylation--- ultrahigh performance liquid chromatography/multiple-reaction monitoring-mass spectrometry (UPLC/MRM-MS) with stable isotope-labeled internal standards for AA quantitation. After validation of the analytical method, the short-term stabilities of 21 AAs in the DBS samples were measured under the various testing conditions.

## Materials and Methods

### CHEMICALS, STANDARD SUBSTANCES AND INTERNAL STANDARDS

Methanol, acetonitrile, isopropanol, water, trifluoracetic acid (TFA), and formic acid were LC/MS grade and were obtained from Sigma-Aldrich (St. Louise, MO). Dansyl chloride was purchased from TCI America (Portland, OR). Authentic compounds of the 21 AAs, including *L*-alanine (*Ala*), *L*-arginine (*Arg*).HCl, *L*-aspartic acid (*Asp*), *L*-asparagine (*Asn*), *L*-glutamine (*Gln*), *L*-glutamic acid (*Glu*), glycine (*Gly*), *L*-histidine (*His*).HCl.H_2_O, *L*-isoleucine (*Ile*), *L*-leucine (*Leu*), *L*-lysine (*Lys*).HCl, *L*-methionine (*Met*), *L*-phenylalanine (*Phe*), *L*-proline (*Pro*), *L*-serine (*Ser*), *L*-threonine (*Thr*), *L*-tryptophan (*Trp*), *L*-tyrosine (*Tyr*) and *L*-valine (*Val*), all of which were from Sigma-Aldrich, together with *L*-ornithine (*Orn*).2HCl and *DL*-citrulline (*Cit*), both of which were from TCI America, were used as the standard substances. Their corresponding ^13^C- or deuterium (D)-labeled analogues, including *L*-ornitinie-^13^C_6_ and *L*-threonine-^13^C_4_ from Sigma-Aldrich; *DL*-Alanine-D_3_, *L*-aspartic-D_3_ acid, *L*-citrulline-D_7_, *DL*-glutamic-D_3_ acid, *L*-histidine-D_6_.HCl.H_2_O, *L*-leucine-D_3_, *DL*-lysine-D_4_.2HCl, *DL*-methionine-D_3_, *L*-proline-D_3_, *L*-serine-D_3_, *L*-tyrosine-D_3_, *D*-tryptophan-D_5_ and *D*-valine-D_8_ from CDN Isotopes Inc. (Point-Claire, QC, Canada); arginine-^13^C_6_, glycine-D_2_-^15^N and *L*-isoleucine-^13^C_6_ from Cambridge Isotope Laboratories Inc. (Tewksbury, MA) as well as *L*-glutamine-^13^C_5_, *L*-phenylalanine-^13^C_6_, and *L*-asparagine-D_3_ from Toronto Research Chemicals Inc. (Toronto, ON, Canada), were used as stable isotope-labeled internal standards (IS) for quantitation.

### WHOLE BLOOD AND DBS PREPARATION

Whatman 903 Protein Saver (abbreviated here as 903; lot No. 7027015 W142), Whatman FTA DMPK-C (abbreviated here as DMPK-C; lot No. ET70110315), and PerkinElmer 226 Spot Saver (abbreviated here as 226; lot No. 101535/1005002) collection cards were used for preparation of the DBS samples used in this study. The absorbing matrix on the 903 and DMPK-C cards was cellulose filter paper, while the 226 cards had cotton filter paper as the blood blotting medium. A single batch of whole blood, which had been collected and pooled from healthy female adult donators, was obtained from BioreclamationIVT (Westbury, NY). We measured the blood hematocrit as ca. 49% before use. The blood was shipped from the supplier at low temperature on ice packs within 48 hours after blood collection, and the necessary biological safety tests had been performed. The whole blood was agitated with a magnetic stirrer at 500 rpm on ice during the whole process of DBS preparation, which involved precisely pipetting and carefully spotting 30-μL aliquots onto the cellulose or cotton linter filter paper of the collection cards. The spotted blood was allowed to complete dry for 4 h in a fume hood at room temperature before the cards were placed into plastic bags, zipped closed, and stored at -80 °C. The use of human blood and DBSs in this study was approved by the research ethics committee at University of Victoria.

### STABILITY TESTING

Four sets of stability-testing experiments were performed. These were ***(1)*** sunlight exposure by placing the DBS cards under direct sunlight for 4 h (the air temperature during the exposure was 21 °C on the day); ***(2)*** three cycles of two-temperature storage by sealing the DBS cards in individual plastic bags containing two small packs of Rubin(tm) indicating silica gel (6g/bag) (Sigma-Aldrich) and placing the bags at -20 °C for 2 days, then at 40 °C for 2 days and back to -20 °C for 2 more days; ***(3)*** storage of DBS cards at four temperatures by placing the samples in the dark at -20 °C (freezer), 4 °C (refrigerator), 25 °C (oven), and 40 °C (oven) for 2, 6, 15,or 30 days, respectively, under laboratory humidity conditions (ca. 38% ± 1% RH); and ***(4)*** storage at two temperatures (25 and 40 °C) at 75% ±1% RH by placing the DBS cards inside a BTL-433 ESPEC humidity chamber (Hudsonville, MI) maintained at 25 or 40°C, respectively, for 2, 6, 15, or 30 days. At each time point during the testing, the DBS samples were collected and processed as soon as possible for the AA quantitation using the optimized and validated analytical method (described below). The number (n) of sample replicates for the AA quantitation was 4. For each round of the AA quantitation, the DBS samples that had been stored at -80 °C were used as controls and the measured concentrations of individual AAs in them were used to normalize the concentrations of the corresponding AAs determined in the spots that were subjected to the stability testing conditions.

### OPTIMIZATION OF SAMPLE PREPARATION FOR AA EXTRACTION

The DBS samples were punched out of the collection cards as whole spots with a 12-mm I.D. handheld stainless steel hole-punch with the aid of a small hammer, and the whole spots were then cut into small pieces and put into 2-mL capless Eppendorf tubes. The AAs were extracted from the DBSs either in a single step or in two steps, using different solvents. For single-step extractions, 1 mL of methanol, acetonitrile, methanol:acetonitrile (1:1, v/v), 80% aqueous acetonitrile, or 80% aqueous methanol:acetonitrile (1:1, v/v), with or without 0.01% to 0.05% TFA (v/v) in each solvent, was added to the sample tubes. The tubes were capped and violently shaken on a digital mixer at 3,000 rpm for 15 s to 2 min, sonicated in an ice-water batch for 2 to 10 min, and then centrifuged at 21,000 x g and at 10 °C for 10 min in an Eppendorf 5420 R centrifuge. For two-step extractions, 200 μL of water or 0.01% to 0.05% TFA in water was added to each tube, followed by the vortex-mixing for 15 s at 3,000 rpm. After a 5-s spin-down in the centrifuge, 800 μL of acetonitrile or methanol:acetonitrile (1:1. v/v), with or without 0.01% to 0.05% TFA in each solvent was added to each tube. The tubes were vortexed again for 15 s, followed by sonication for 2 to 10 min, before centrifugation at 21,000 x g for 10 min. After the one-step or two-step extractions and clarification by centrifugation, precise 50-μL aliquots of the supernatants were removed and transferred to another set of Eppendorf tubes, and dried in a speed-vacuum concentrator. The residues were for dansylation and UPLC/MRM-MS, as described below. The extraction efficiency was expressed as the % of the average peak area of an AA measured in extracts of triplicate spots from each DBS set, relative to the average peak area measured, in the same extraction experiment, from 30-μL aliquots (n=3) of the liquid whole blood that had been stored at -80 °C.

### UPLC/MRM-MS

The LC/MS system was an Agilent 1290 liquid chromatograph coupled to an Agilent 6490 triple-quadrupole mass spectrometer (Santa Clara, CA), which was equipped with an ESI ion source and operated in the positive-ion MRM mode. The Q1 to Q3 ion transitions (2 transitions per analyte) and corresponding collision energies of dansylated AAs for MRM-MS were optimized using Agilent’s *MassHunter Optimizer* while infusing a standard solution of each derivatized AA. An Agilent Eclipse Plus RRHD C_18_ column (2.1 mm I.D. x 150 mm, 1.8 μm) was used for the LC separation, with 0.1% (v/v) formic acid in water (solvent A) and 0.1% formic acid in acetonitrile:isopropanol (1:1, v/v) (solvent B) as the mobile phases for binary-solvent gradient elution, with an optimized gradient of 20% to 50% B in 12 min and 50% to 100% B in 4 min and a flow rate of 0.325 mL/min. The column was then washed with 100% B for 1 min, followed by a 3-min column-equilibration between injections. The column temperature was held at 50 °C and the autosampler temperature was maintained at 5 °C.

### STANDARD SOLUTIONS, DERIVATIZATION, AND QUANTITATION

Stock solutions of the 21 AAs were prepared individually by dissolving the standard substances with 50% aqueous acetonitrile to produce the same concentration of 4.2 mM. The stock solutions were mixed at an equal volume to make a concentrated standard solution, and a serial dilution of this solution was prepared at a ratio of 1 to 4 (v/v) with 50% aqueous acetonitrile to generate working standard solutions with a concentration range of 50 μM to 3 nM for each AA. For preparing the calibration curves of 21 AAs for quantitation, 50 μL of each working standard solution was added, followed by 50 μL of the IS solution that contained the ^13^C- or deuterium (D)-labeled IS compounds in a concentration range of 0.5 to 5 μM for individual AAs, depending on their endogenous concentrations, in 50% aqueous acetonitrile, 100 μL of 40 mM dansyl chloride in acetonitrile, and 50 μL of 100 mM borate buffer (pH9.2) (in this order). For derivatization of the samples, the dried residue from 50 μL of the clear supernatant extracted from each dried blood spot (as described in the “*Optimization of sample preparation for AA extraction*” section), was resuspended in 50 μL of 50% aqueous acetonitrile, followed by sequential addition of the IS solution, the dansyl chloride solution, and the borate buffer in the same order as was used for the standard solutions. The mixtures were allowed to react at 40 °C for 40 min. After the reaction, 750 μL of 50% aqueous methanol was added. After vortex mixing and centrifugation at 21,000 x g for 10 min, 10-μL aliquots of the clear supernatants were injected for UPLC/MRM-MS. The concentrations of the 21 AAs in DBSs were calculated from the linear-regression calibration curves of individual AAs with IS calibration. All of the data was recorded and processed with the Agilent’s *MassHunter* software suite. Analysis of variance (ANOVA) and student’s t-test statistics were performed using the SYSTAT software suite (San Jose, CA).

### VALIDATION OF QUANTITATION

The intra-day precision was determined as the relative standard deviations (RSDs) obtained when the AA quantitation for each DBS set was performed every 3 h for a total of 6 times (n=6) within the same day; the inter-day precision was determined as the RSDs obtained when the AA quantitation was performed on the same DBS sets for 5 continuous days. The quantitation accuracy was determined from the standard-substance spiking recoveries tested at three spiking levels, equivalent to 100%, 250%, and 500% of the endogenous concentrations of each AA measured in the DBS samples. The recoveries were calculated as (total concentration - endogenous concentration)/spiked-in concentration × 100%. The number (n) of sample replicates for each spiking level was 5.

## Results

### EXTRACTION OF AAs

AAs in DBS specimens are usually extracted with methanol, then butylated and analyzed by FIA-MS/MS with neutral loss scanning to provide rapid analysis for IEM screening (16, 18, 19). However, this analytical technique can only be used for measurements of some of the AAs (19). For example, *Asp* and *Glu* could not be measured due to the signal overlap with *Asn* and *Gln* in the infusion-MS/MS experiment (16). To provide precise and accurate measurements of all of the 21 AAs for evaluation of their short-term stabilities in DBSs, we chose UPLC/MRM-MS with stable-isotope labeled internal standards to provide a separation step and more accurate quantitation. First, we revisited the “standard” extraction procedure by comparing methanol with two other solvent systems, i.e., acetonitrile, and methanol:acetonitrile (1:1, v/v), to extract AAs from the DBS samples on the different collection cards. Mixed methanol:acetonitrile has been reported to be superior to methanol or acetonitrile alone for extractions of endogenous metabolites in blood plasma in LC/MS based metabolic profiling (20). Measurements of the 21 AAs from the extracts using these three solvents and the optimized dansylation-UPLC/MRM-MS method (described below), showed that complete extraction of any of the 21 AAs from the DBSs were difficult to achieve with any of them. Specifically, the basic *Arg, Lys, Cit, Orn*, and the weakly basic *His*, the aromatic *Tyr, Phe* and *Trp*, and the amide side chain-containing *Asn* and *Gln*, had significantly lower extraction efficiencies than the other AAs, even with extended periods of sample vortex-mixing (15 s to 2 min) and sonication (2 to 10 min). The use of methanol, acetonitrile, or methanol:acetonitrile (1:1) produced only average extraction efficiencies of ca. 50%, 57%, and 74%, respectively, for the 21 AAs from the three sets of DBS samples. Considering the basicity of the basic AAs and the dipolar nature of AAs, we next evaluated extractions done with the three solvents containing 0.01% to 0.05% TFA (v:v), and their 80% aqueous solutions (except for 80% aqueous methanol, which did not precipitate the protein very well during the extraction procedure), with or without 0.01% to 0.05% TFA in each solvent. The inclusion of TFA as the acidic modifier and the formation of “neutral” ion pairs between TFA and AAs in the extracting solvents significantly improved the extraction efficiencies for all of those AAs that had shown lower-extraction efficiencies when the TFA-free solvents were used. For example, the use of methanol, 80% acetonitrile, or 80% methanol:acetonitrile (1:1), containing 0.02% TFA in each solvent, gave average efficiencies of 72%, 79%, and 88% extraction for the 21 AAs. However, none of the TFA-containing solvents produced satisfactory extractions for the majority of 21 AAs in a single-step extraction procedure. We hypothesize that this might be because a pure or high-percentage (i.e., 80%) of organic solvent, was unable to completely resuspend the whole blood after drying-down on the cellulose or cotton filter paper even when a low % of TFA was included in the extraction solvent. Thus, free AAs might not be completely released during these extractions.

We then evaluated two-step extractions to completely resuspend the whole blood from the filter paper, using 200 μL of water containing 5 concentrations of TFA (0.01%, 0.02%, 0.03%, 0.04%, and 0.05%) as the first step and then using 800 μL of acetonitrile or methanol:acetonitrile (1:1) containing the same concentrations of TFA as used in the first step (i.e., 0.01%, 0.02%, 0.03%, 0.04%, and 0.05%), as a second step, for a total of 10 combinations. These two-step extractions were found to be better than the single-step extractions -- the average extraction efficiencies were 91% and 99%, respectively, for the 21 AAs, when acetonitrile or methanol:acetonitrile (1:1), containing 0.02% TFA in each solvent, were used in the second-step extractions. The highest yield and reproducible extractions were from the two-step extraction using the combination of 200 μL of water containing ≥0.02%TFA and 800 μL of methanol:acetonitrile (1:1) containing ≥0.02% TFA for all the three sets of DBSs. Also, the concentrations of TFA in the range of 0.02% to 0.05% did not generate any significant differences in the extraction efficiencies. Supplemental Information Fig. S1 shows a comparison of the extractions for the five basic AAs (*Arg, Lys, Cit, Orn* and *His*), the three aromatic AAs, and *Asn, Gln, Asp*, and *Glu* from the three sets of the DBS samples using either A) 200 μL of 0.02% TFA in water and 800 μL of acetonitrile containing 0.02% TFA or B) methanol:acetonitrile (1:1) containing 0.02% TFA, for the two-step extractions, or C) using methanol:0.02% TFA for the single-step extractions. Based on these comparisons, the optimized extraction procedure for all the subsequent experiments in this study was done using two-step extractions with the combination of 200μL of 0.02% TFA in water and 800 μLof methanol:acetonitrile:TFA (1:1) containing 0.02% TFA.

### AA QUANTITATION BY DANSYLATION-UPLC/MRM-MS

DBS specimens have complicated biological matrices in comparison with blood serum or plasma, as DBS samples not only contain most components existing in serum or plasma, but also include dried blood cells, etc. The blood-spotted filter paper may also adversely affect accurate measurements of specific molecular targets. Therefore, to provide accurate measurements of all the 21 AAs, we combined the optimized extraction procedure with chemical derivatization-UPLC/MRM-MS with dansyl chloride, a commonly used derivatizing reagent for amino group-containing compounds (21), for pre-column reaction with the 21 AAs. Based on the dansylation reaction conditions described in previous studies on the LC/MRM-MS quantitation of AAs (21-23), the reaction temperatures (30, 40, 50, and 60 °C), the reaction time periods (10, 25, 40, 55, 70, 85 min), the %s of acetonitrile (30%, 45%, 60%, and 75%), and the pH of the reaction medium (in a range of pH 8.5 to 10.5), as well as post-reaction treatments,were re-examined. We determined the optimal reaction conditions to be 40 °C for 40 min in 60% aqueous acetonitrile and at pH 9.2 in the borate buffer. Under these reaction conditions, 17 of the 21 AAs were singly derivatized with dansyl chloride, while the other 4 AAs (*Tyr*, *His, Orn* and *Lys*) were doubly dansylated with no partially derivatized products, as determined by UPLC/MRM-MS using the ion transitions for the singly-derivatized products (data not shown). Supplemental Information Fig.S2 shows a representative UPLC/MRM-MS chromatogram of the 21 dansylated AAs separated on the C_18_ column with 0.1% formic acid in water (solvent A) and 0.1% formic acid in acetonitrile:isopropanol (1:1, v:v) (solvent B) as the binary solvents for gradient elution. As shown, isomeric Leu and Ile were well separated, and *Asn* and *Gln* were also well separated from *Asp* and *Glu*, respectively, which eliminates any possible signal overlap between them. For the post-reaction treatment, we diluted the reactant solutions with 50% methanol in a 1:4 (v:v) ratio, instead of quenching the reaction with a high-concentration ammonia solution (21). When this 50% methanol procedure was used, the ratios of the peak areas of the unlabeled endogenous forms of the 21 derivatized AAs versus their stable isotopically-labeled IS compounds (i.e., As/Ai) -- as measured from sample solutions and standard solutions --were stable at 5 °C in the autosampler for at least 90 h. The RSDs of the As/Ai ratios measured within 90 h for all the 21 AAs were ≤ 6.3% in all cases. Supplemental Information Fig. S3 shows the measured As/Ai ratios of eight representative AAs from a derivatized sample solution from each of the three sets of DBSs, indicating the high stabilities of the AA derivatives. Also, no precipitates resulting from excess dansyl chloride or from any AA derivative were seen in the 50% methanol diluted sample solutions during the UPLC/MRM-MS analyses of large batches of the samples.

Dansylation of AAs and amines, like many other amino group-specific chemical derivatization approaches, needs to be carried out in a basic reaction medium, with a pH of ca.8.5-10.5, in order to drive the reactions to completion (23). Based on our observations during preliminary testing, small portions of *Asn* and *Gln* can deamidate to form *Asp* and *Glu*, respectively, in this pH range. Also, a small amount (≤ 8%) of *Glu* underwent intra-molecular cyclization to form the less polar pyroglutamic acid by loss of a water molecule from *Glu*, as observed by using a hydrophilic interaction LC/MS method (data not shown). These side reactions were independent of the presence of dansyl chloride. To compensate for the influences of these side reactions at the basic pH, as well as to compensate for ionization suppression from the DBS sample matrices during LC/MRM-MS runs, we found it necessary to use^13^C- or D-labeled analogues of the 21 AAs as internal standards to achieve accurate quantitation. Supplemental Information Table S1 shows the endogenous concentrations of the 21 AAs measured in the control samples for each of three sets of DBSs. As part of the validation of the quantitation --as well as to test whether the cellulose or cotton filter paper had adverse effects on the AA quantitation --we followed the US-FDA Guideline for Bioanalysis Method Validation for Industry (24) and evaluated the assay precision by measuring the intra-day and inter-day RSDs, and the assay accuracies by determining the standard-substance spiking recoveries at spiking levels of 100%, 250%,and 500% of the endogenous concentrations for each AA in each of the three sets of DBSs. Supplemental Information Table S2 summarizes the results of these precision and accuracy measurements. As shown, all of the intra-day and inter-day RSDs were ≤7.9% (n=6) and ≤6.3% (n=5), in all cases. The measured recoveries for the 21 AAs ranged from 88.4% (RSD =3.6%, n=5) for *Asp* to 112.8% (RSD=4.1%, n=5) for *Met*, at the low, medium, and high spiking levels, which is indicative of good quantitation accuracy. Student’s t-test statistics indicated that there were no significant differences at P < 0.05 in the measured precision and accuracy of the quantitation among the three sets of DBSs. This validated method was then used to determine the concentrations of the 21 AAs in the DBS samples under the different environmental conditions tested in this study.

### STABILITY OF AAs AFTER 4-HOUR EXPOSURE TO SUNLIGHT

Fig. 1 is a bar graph showing the % concentrations and standard deviations (SDs) of the 21 AAs, as determined from four spots (n=4) per DBS set, which had been exposed to direct sunlight irradiation for 4 hours. The concentrations of each AA in the samples have been normalized to those measured in the controls for each set, which were stored at -80 °C. All 21 AAs were stable in the three sets of DBS samples during a 4-h exposure to sunlight. The only AA that showed more than 10% concentration decreases in the DBS samples on two of the three types of collection cards was *Met*, whose concentration decreased by 15.5% and 10.9% on the filter paper of the 903 and 226 cards, respectively. The degradation of *Met* is probably due to the existence of an oxidization-susceptible S-methyl thioether group in its side chain. The average concentrations ± RSDs for all of the AAs (n=21) were 94.4% ± 3.2%, 96.8% ± 2.5%, and 99.4% ± 3.6% for the 903, DMPK-C, and 226 cards, respectively, which implied nearly equal stabilities of the AAs on the three sets of collection cards.

**Fig. 1.**
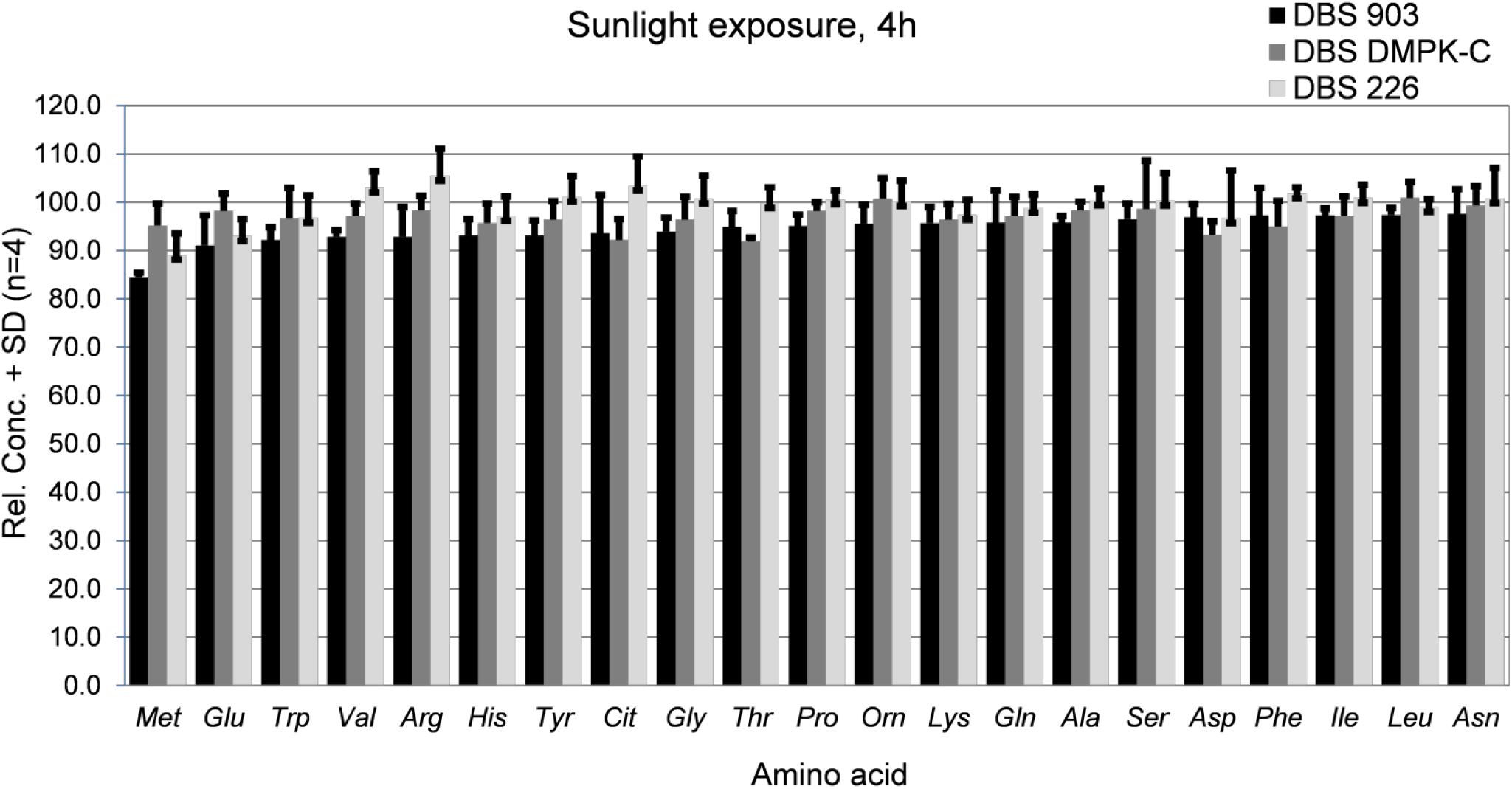
Relative concentrations and SDs (n=4) of 21 AAs measured in DBSs on three sets of collection cards (226, 903 and DMPK-C) after 4-h sunlight exposure.

### STABILITY OF AAs AFTER THREE CYCLES OF TWO-TEMPERATURE STORAGE

To test the effect of temperature transitions which could be encountered during the sample transportation and storage in global health applications, especially for DBS specimens collected in tropical sampling locations, on the stabilities of the different AAs, the three sets of DBSs were stored in desiccated bags and underwent three cycles of two-temperature storage for a total of 6 days: -20 °C for 2 days, 40 °C for 2 days and back to -20 °C for 2 additional days. The bar graphs in Fig. 2 show the relative concentrations and SDs (n=4) of the 21 AAs measured in the three sets of DBS samples following this temperature cycle. As can be seen from this figure, all of the 21 AAs degraded, to a greater or lesser extent, during the temperature cycle under the drying silica gel-reduced humidity. *His* was the least stable AA -- its concentration decreased by 30.7%, 28.4%, and 39.7%, on the 226, 903, and DMPK-C cards, respectively. Another AA that also showed a relatively high extent of degradation was *Met*, whose concentration decreased by 25.2%, 27.2%, and 29.5%, respectively, in the three sets of DBS samples. *Gln* showed average concentration decreases of 14.2%, 19.9%, and 15.7%, respectively, in the three sets of DBS samples. In addition to the above, *Thr*, *Orn*, and *Ser* showed average concentration decreases of 14.3%, 11.6%, and 11.5%, respectively, in the DBS samples on the three types of collection cards. The other 15 AAs had average concentration decreases of <10% on the three types of cards, indicating the good stabilities of these AAs.

**Fig. 2.**
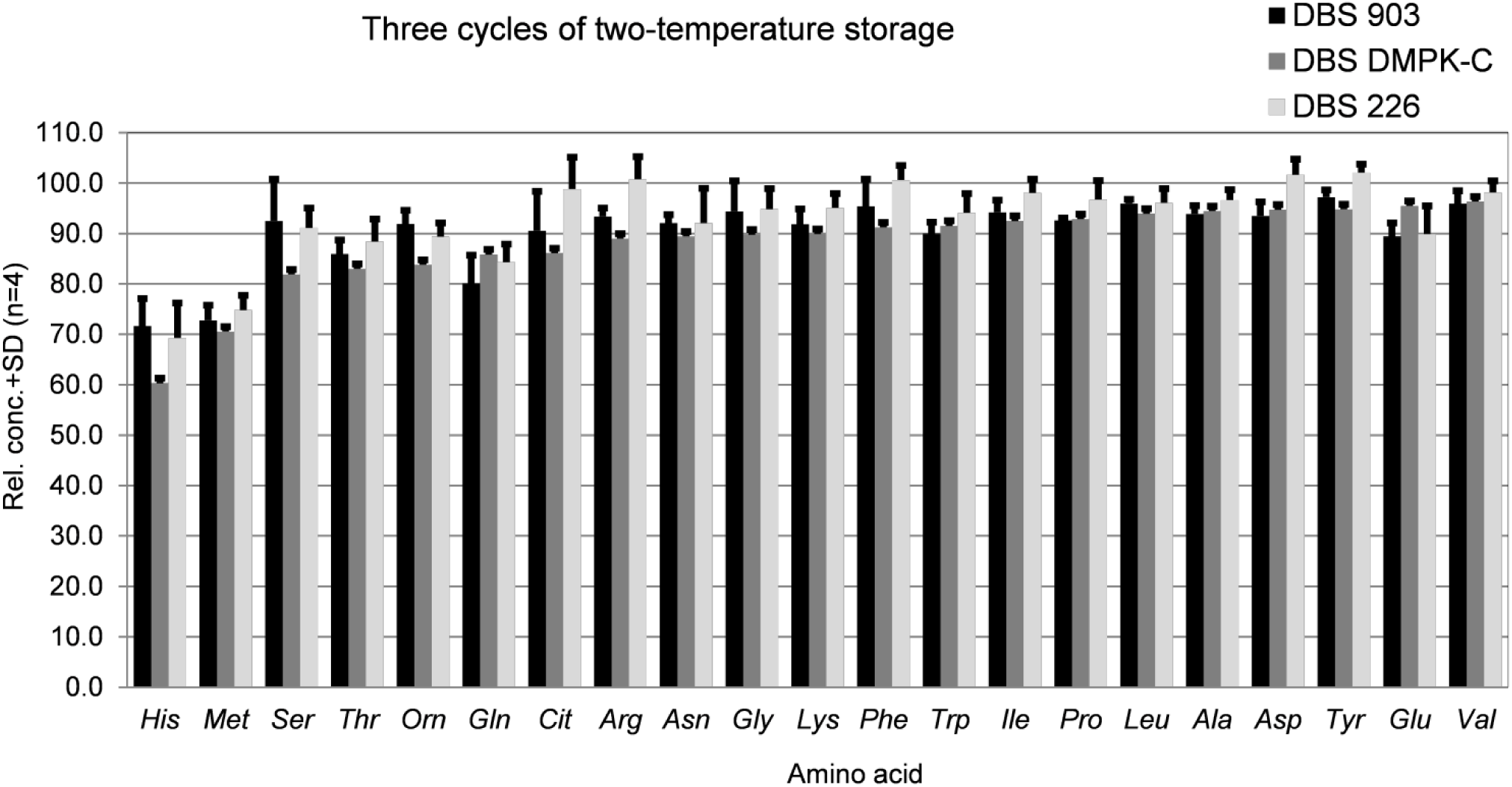
Relative concentrations and SDs (n=4) of 21 AAs measured in DBSs on three sets of collection cards (226, 903 and DMPK-C) after three cycles of two-temperature storage (-20 °C for 2 days, 40 °C for 2 days and -20 °C for 2 additional days) in desiccated bags.

The average relative concentrations (as %s ±SDs) of the 21 AAs (n=21) from the 226, 903, and DMPK-C cards were 93.0% ±9.1%, 90.2%±7.8%, and 88.0%±9.9%, respectively, indicating quite comparable stabilities of the AAs on the three sets of DBS samples. Paired and one-tailed T-tests indicated that there was a slightly statistical significance at P < 0.01 (the 99% confidence level) in the stabilities of these 21 AAs, on the whole, between the 266 card and the DMPK-C or 903 cards, but not between the 903 card and the DMPK-C card. However, further studies are needed to determine whether the statistical difference in the AA stabilities between the 226 cotton filter paper card and the other two sets of cellulose filter paper cards can be attributed to the differences in the blood absorbing matrices. This stability testing demonstrated that the temperature transitions could affect the concentrations of certain AAs in DBSs, even under conditions of reduced humidity, and implied that exposure of DBSs to some extreme environmental stress, e.g., the 40 °C storage temperature, should be avoided.

### STABILITY OF AAs IN DBS AT DIFFERENT STORAGE TEMPERATURES

To further investigate the stabilities of various AAs in DBSs, as affected by sample storage at different temperatures, the three sets of DBS samples were stored at four temperatures (-20 °C, 4 °C, 25 °C, or 40 °C) in the dark and under laboratory humidity (ca. 38% ±2% RH as monitored throughout the experiments) for 2, 6, 15, and 30 days, respectively. Supplemental Information Fig. S4 shows bar graphs indicating the relative concentrations and SDs (n=4) of the 21 AAs measured in each sample set. *Ala, Asn, Cit, Gln, Ile, Leu, Pro, Met*, and *Trp* did not show significant concentration decreases -- ≤10% at -20 °C, 4 °C, and 25 °C for 30 days in all the three sets of DBSs. *Gly, His, Lys, Orn*, *Thr*, and *Tyr* were stable in all of the DBSs at -20 °C and 4 °C for 30 days, but at 25 °C these AAs showed concentration decreases starting from 6 days of sample storage, and continuing for the entire 30-day period at 40 °C in a time-course dependent manner. *His* was the least stable of all of the tested AAs in this experiment. Its concentrations decreased at 25 °C and 40 °C in a time-course and temperature-dependent way. At 40 °C, its concentrations dropped by 38.2%, 57.8%, and 68.1% on the cotton paper of the 226 card or the cellulose paper of the 903 and DMPK-C cards, respectively, at 30 days. *Lys* showed good stability at -20 °C for 30 days and at 4 °C for up to 15 days. It showed >10% degradation at 4 °C in 30 days. The degradation of *Lys* increased at higher storage temperatures, showing a decrease of 22.3%, 32.5%, and 34.2% on the filter paper of the 226, 903, and DMPK-C cards, respectively, when stored at 40 °C for 30 days.

The two acidic AAs *(Asp* and *Glu)* showed concentration trends that were different from the other AAs, as seen in Supplemental Information Fig. S4. At -20 °C and 4 °C for 30 days, and at 25 °C for 6 days, *Asp* underwent concentration <u>increases</u> in all three sets of DBSs, with an increase of up to 26.2% on the 226 card. *Glu* showed concentration increases on the 226 and DMPK-C cards, with an increase of 19.3% on the 226 card. The source of these concentration increases for these two acidic AAs was not determined. One explanation might be that the increases could result from deamidation of *Asn* and *Gln*, though no significant concentration changes were observed for free *Asn* and *Gln* in the same DBS samples stored at the low temperatures. At 25 °C for 15 and 30 days, and at 40 °C over 30 days, the two acidic AAs displayed time-course-related decreases in concentration, and the concentrations dropped more rapidly at 40 °C than at 25 °C. This indicates that self-degradation was the major mechanism that contributed to their concentration changes at the higher temperatures.

To evaluate the differences in the stabilities of different AAs in the three sets of DBSs on the cellulose or cotton filter paper, one-way ANOVA statistical analysis was performed on the concentrations of the 21 AAs measured in the DBSs on each of the three card types, which were tested at the four temperatures over 30 days. At a significance level of P < 0.01, no significant difference among the three sets of DBSs were observed for the samples stored at -20, 4, or 25 °C over 30 days, indicating comparable stabilities of the AAs on these cards at these temperatures. However, using the same significance value (P < 0.01) cutoff, there was a statistical difference (P = 0.005< 0.01) in the measured concentrations from the samples stored at 40 °C over 30 days among the three sets of DBSs. The average concentrations of the 21 AAs ± their SDs (n=21) on the 226, 903, and DMPK-C cards were 93.6% ±14.9%, 86.3% ±15.0, and 82.7% ±16.1%. Paired and one-tailed student’s tests between any two groups indicated that the statistical significance came from the concentration difference between the 226 card and the 903 card or the DMPK-C card, but not between the 903 and DMPK-C card. This observation, together with the similar observation from the three-cycle temperature-related stability testing in which the samples had been exposed to the same high temperature (40 °C) for 2 days, as described above, implied that the two types of blood absorbing matrices (i.e., cotton on the 226 card and cellulose on the 903 and DMPK-C cards) might have a slightly different effect on the chemical stabilities of the AAs in the DBSs. However, more experiments need to be performed to confirm these observations.

### STABILITY OF AAs IN DBS UNDER HIGH HUMIDITY

To test the influence of high humidity on the stabilities of different AAs in DBSs, the three sets of DBS samples were stored at 25 °C and 40 °C at the same high humidity level (75%±1% RH) for 2, 6, 15, or 30 days, respectively. Supplemental Information Fig. S5 shows the bar graphs for the measured % concentrations and SDs (n=4) of 21 AAs in each set of the DBSs, relative to the concentrations in the corresponding controls. The trends of these concentration changes over time in three sets of DBS at 25 °C and 40 °C are shown in Fig. 3.

**Fig. 3.**
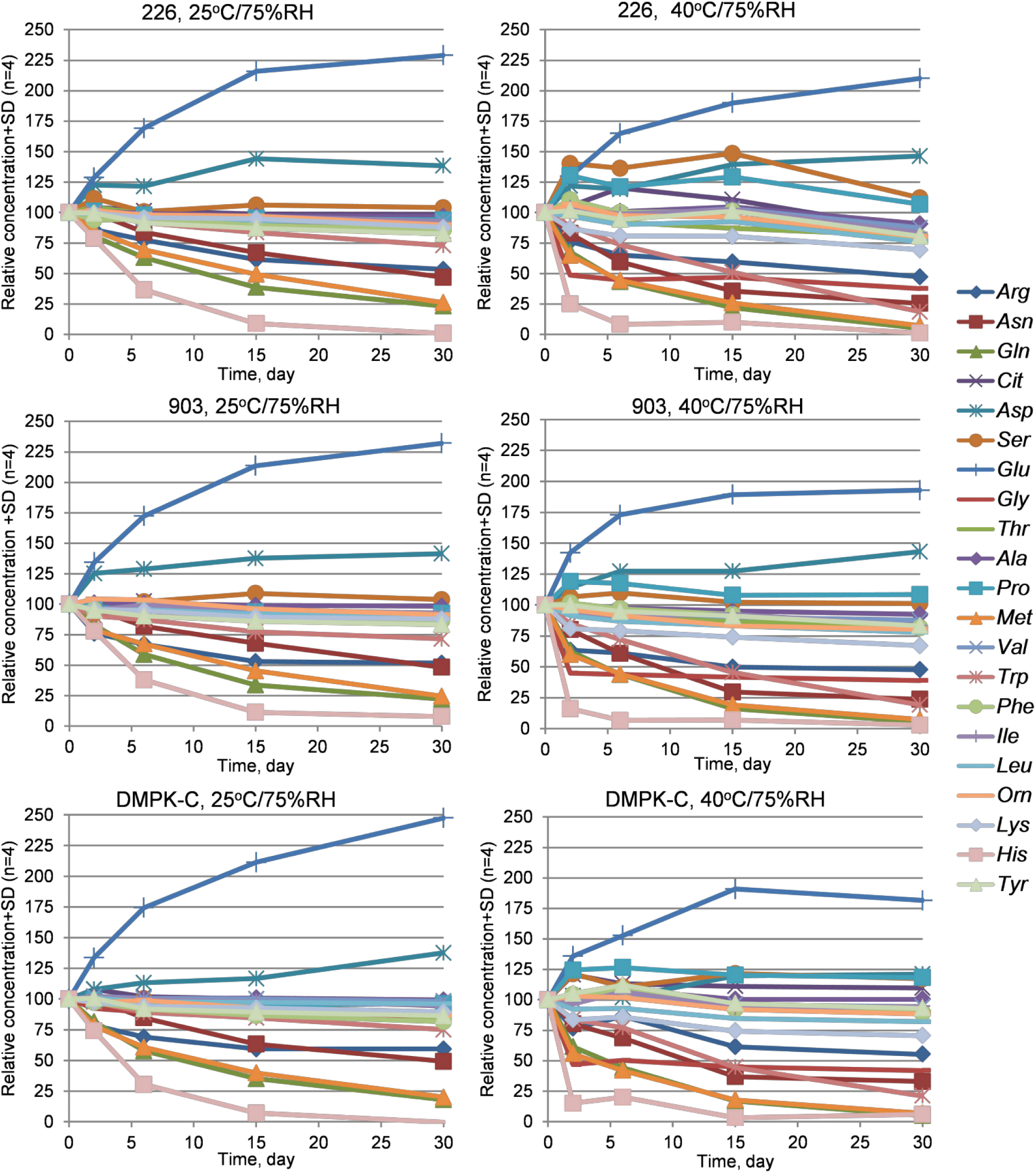
Concentration changes of 21 AAs over time in DBSs on three sets of collection cards (226, 903, and DMPK-C) at storage temperatures of 25 °C and 40 °C and at 75% RH.

Among the 21 analytes, 19 AAs, except *Glu* and *Asp*, showed time-course concentration decreases when stored at 75% RH at both temperatures. The concentration decreases were larger at 40 °C under the high humidity for most of these 19 AAs than at 25 °C, for each of the three DBS sets. The order of lower to higher percent changes in the normalized concentrations is *Ala<Ile<Leu<Val<Pro<Tyr<Thr<Cit*<*Lys, all of which* showed concentration decreases of ≤ 20% at 40 °C, followed by *Phe<Orn<Gly<Ser*, which concentrations decreased between 20% and 40% in the three sets of DBSs, after 30 days of storage. The concentrations of *Arg* decreased by 32%-45% in the DBSs stored at 40 °C for 30 days. For the remaining AAs, the concentrations of *Asn, Trp, Met*, and *Gln* decreased by 70-72%, 80-81%, 87-90%, and 93-95%, respectively, in the three sets of DBSs stored at 40 °C, and their degradation at 40 °C was much faster than at 25 °C, as shown in Fig. 3. The concentration of *His*, which was found to be the *least* stable of the 21 AAs in the above two storage temperature-related stability testing experiments, decreased by >97% at both 25 °C and 40 °C at 75% RH, showing that it was the most unstable AA under the high-RH testing conditions as well.

In contrast to the above-mentioned 19 AAs, the two acidic AAs (*Asp* and *Glu*), showed temperature-dependent concentration increases during the 30-day time course of the experiment. The concentrations of *Asp* and *Glu* <u>increased</u> at 30 days by 38%-43% and by 120%-144%, respectively, in the three sets of DBSs at 25 °C, and by 20%-41%, and by 80%-112%, respectively, at 40 °C. As mentioned above, these concentration increases might be due to the contribution from the deamidation of *Asn* and *Gln* and/or protein hydrolysis under the high humidity conditions. Comparing the concentrations of different AAs in the DBSs stored at 25 °C and 40 °C under laboratory humidity (ca. 38% RH), the concentration changes of all the AAs were accelerated by higher humidity.

To assess if there are significant differences in the AA stabilities among the three sets of DBSs at 75% RH, one-way ANOVA and student’s T-tests were applied to the measured concentrations of the 21 AAs in the three sets of DBSs stored at 25 °C and 40 °C, respectively, at 75% RH, for 30 days. No statistical significance was observed among the three sample groups or between any two groups for each temperature, which indicated the comparable stabilities of these AAs on the cellulose and cotton filter paper with regard to the RH.

## Discussion

To prepare DBSs for quantitative analysis, the whole blood was stirred before spotting, and the entire blood spot -- instead of a partially cut disk --was used as the starting material for the subsequent AA extractions. These steps helped to minimize the analytical variations resulting from the DBS preparation and partial-spot sampling. With the combination of the dansylation-UPLC/MRM-MS method, concentrations of the 21 AAs in the DBSs on three types of cellulose or cotton collection cards were precisely and accurately quantitated, which ensured the successful determination of their stabilities in the four testing experiments.

In previous studies on short-term (17) and long-term (16) stabilities, 8 and 10 AAs in DBSs were measured. In both of the studies, *Met* were reported to be the least stable of the targeted AAs. With the expanded panel of 21 AAs involved in this study, and under the well-controlled testing conditions used here, *His*, which was not included in any of the two previous studies (16, 17),was found to be the least stable AA in all the DBSs on three types of collection cards under the temperature and humidity conditions tested, although *Met* was also determined to be one of the less-stable metabolites under all the testing conditions used.

With the 21 AAs involved in this study, *Trp* -- but not the two other aromatic AAs (*Phe* and *Tyr*) -- was found to be less stable under high humidity (Supplemental Information Fig. S5). Comparing the chemical structures among the three aromatic AAs and that of *His*, the facile degradation of *Trp* and *His* is probably due to the chemical instabilities of the heterocyclic indole and imidazoleside chains in their structures, although further investigation is needed to draw a definite conclusion.

Together with *His* and *Trp*, other AAs including *Arg, Lys, Cit, Orn, Met, Asp*, *Glu*, *Asn* and *Gln* also showed larger concentration changes (increases for *Asp* and *Glu* and deceases for the others) in the DBSs under high-humidity conditions. The concentration decreases for the four basic AAs (*Arg, Lys,Cit*, and *Orn*), *Asn*, and *Gln*, seemed to be related to the guanidinium (*Arg*), amino (*Lys* and *Orn*), or amide (*Cit*, *Asn* and *Gln*) functional groups in their side chains. The two acidic AAs, *Asp* and *Glu*, differed from the other AAs in that both showed time-course concentration <u>increases</u> in the DBSs under the same high humidity conditions, which indicates that the contributions to their concentrations from the deamidation of *Asn* and *Gln* and/or protein degradation in the DBSs exceeded the rate of degradation for these two acidic AAs under the conditions tested. Also, at 15 and 30 days, the measured concentrations of *Glu* in the three respective DBS sets at 40 °C were lower than those at 25 °C, which indicated that degradation of this acidic AA at 40 °C proceeded more rapidly than at 25 °C, and that degradation more than compensated for any increase in *Glu* concentration which might have resulted from deamidation of *Asn* and *Gln* and/or protein degradation in the DBSs under the high-humidity conditions.

In this study, all of the stability testing experiments were performed at the endogenous concentrations of the 21 AAs in the DBS specimens, not at different concentration levels At the endogenous level, a statistically significant difference in the concentrations of 21 AAs in the DBSs was observed between the cotton filter paper card (226) and the two types of cellulose filter paper cards (903 and DMPK-C) tested at the high temperature (40 °C). Considering that all the stability testing experiments involved in this study were carried out on a single lot of each type of collection card, and at a single concentration level for each AA, future experiments need to be performed to confirm if the significant differences in the AA stabilities between the 226 card and the 903 or DMPK-C card resulted from the different blood absorbing matrices (i.e., cotton versus cellulose filter paper).

## Conclusions

The short-term stabilities of 21 AAs in DBS specimens on three sets of cellulose or cotton filter paper were evaluated under simulated environmental conditions that would be expected in a global health workflow. The concentrations of these AAs approximated their endogenous concentrations in whole blood, and were measured using our improved sample preparation and the chemical derivatization-UPLC/MRM-MS method. The results indicate that higher sample storage temperatures (25 °C and 40 °C) and high humidity (75% RH) significantly influence the short-term stabilities of some AAs in DBSs on both of cellulose and cotton filter paper. The basic AAs (*Arg, Lys, Cit, and Orn), Met* (which is easily oxidized)*, Trp* (which contains an indole side chain), weakly basic *His* (which contains an imidazole side chain), *Asn* and *Gln* (which are susceptible to deamidation), and *Glu* and *Asp* (which are acidic), are the AAs whose concentrations were the most affected by higher temperatures and high humidity. *His* was found to be the least stable AA under the conditions tested, except for the 4-h sunlight exposure. The results also indicated that there might be some overall differences in the AA stabilities between the cotton filter paper card and the two types of cellulose filter paper cards when DBS samples are stored at high temperature (40 °C), but further investigation need to be done to confirm this observation, considering only a single lot of each type of collection card and only a single, endogenous concentration level for each AA in the whole blood was tested in this study.

In summary, this study provides quantitative data on the short-term stabilities of 21 AAs in DBSs on both cellulose and cotton collection cards, when subjected to the environmental conditions expected in a global health environment. Based on the results of this study, it is recommended that lower-temperature and low-humidity storage conditions are critical for maintaining the stability of some AAs measured from DBS specimens. This study emphasizes the importance of assessing the stability of any clinical or epidemiological biomarker measured from a DBS specimen, particularly if the collection, transportation, and storage might be exposed to uncontrolled environmental conditions.

## Author Contributions

All authors confirmed they have contributed to the intellectual content of this paper and have met the following 3 requirements: (a) significant contributions to the conception and design, acquisition of data, or analysis and interpretation of data; (b) drafting or revising the article for intellectual content; and (c) final approval of the published article.

## Authors’ Disclosures or Potential Conflicts of Interest

No authors declared any potential conflicts of interest.

## Research Funding

Funding to “The Metabolomics Innovation Centre (TMIC)” through the Genome Innovations Network (GIN) from Genome Canada, Genome Alberta and Genome British Columbia for operations (205MET and 7203) and technology development (215MET and MC3T) is acknowledged. This work was also partially supported by the Bill and Melinda Gates Foundation.

## Patents

None declared.

## Acknowledgments

CHB is grateful for support from the Leading Edge Endowment Fund (University of Victoria) and for support from the Segal McGill Chair in Molecular Oncology at McGill University (Montreal, Quebec, Canada). CHB is also grateful for support from the Warren Y. Soper Charitable Trust and the Alvin Segal Family Foundation to the Jewish General Hospital (Montreal, Quebec, Canada).

